# Biochemical, serological and molecular confirmation of *Salmonella* isolates

**DOI:** 10.1101/100917

**Authors:** V. Prasanna Kumar, S. P. Singh, A. K. Upadhyay, Deepak Kumar

## Abstract

Salmonellosis is one of the most frequently reported food-borne diseases world-wide commonly caused by *Salmonella* Typhimurium and *Salmonella* Enteritidis serovars. The present study was undertaken to confirm *Salmonella* isolates by various techniques. All the 94 isolates were confirmed to be S. Typhimurium through various morphological, biochemical, serological and molecular methods.

## Introduction

Salmonellosis is a leading cause of bacterial gastroenteritis world-wide (**Majowicz *et al.*, 2010**), which may vary in severity from self-limiting infections to acute disease. Humans generally acquire *Salmonella* infection through the consumption of contaminated foods such as beef, chicken, eggs, milk and their products and this may be fatal to the young, the old or the immuno-compromised patients (**Gibbons, 1980**). In animals also, some of *Salmonella* infections are severe and fatal due to the involvement of host specific serotypes.

The occurrence of diarrheal diseases due to *Salmonella* serotypes is a global phenomenon with hospitalization of 93.8 million human cases and 155,000 deaths each year (**Hendriksen *et al.*, 2011**). Salmonellosis is largely caused by *S.* Typhimurium and *S.* Enteritidis, where *S.* Typhimurium infection has been most frequently found associated with consumption of foods of poultry and bovine origin (**Capuano *et al.*, 2013**). Pathogenicity of *Salmonella* is multifactorial and is controlled by group of genes in *Salmonella* pathogenic islands (SPI). It can also cause severe invasive disease in HIV-infected, malaria-infected and malnourished children (**Ao *et al.*, 2015**), which require antibiotic treatment with extended-spectrum cephalosporins, especially to treat children (**Threlfall, 2002**).

## Materials and methods

### Bacterial strains

A total of 94 *Salmonella* suspected isolates of diverse origin were procured from different parts of North Indian states. These bacterial strains were tested for their purity, morphology and other biochemical characteristics.

### Revival of the bacterial isolates

*Salmonella* cultures received from various places were revived using Brain Heart Infusion (BHI) broth and overnight incubation at 37°C. The broth cultures were streaked on brilliant green agar (BGA). All the isolates were revived, tested for their purity, morphology and biochemical characteristics as per standard protocol described in Laboratory Manual for the Isolation and Identification of Food-borne Pathogens (**Agarwal *et al.*, 2003**). The revived isolates were maintained on nutrient agar slants at 4°C and the same isolates were also stored at - 80°C in 15% glycerol stock for long term preservation.

### Biochemical characterization

All *Salmonella* isolates were subjected to cultural characterization. Moderately large, moist, smooth and colourless with pink background on brilliant green agar (BGA) were picked up and confirmed biochemically as per the protocol described by **Ewing (1986)**. Biochemical characterization involved three tests namely triple sugar iron (TSI) test, urease test and citrate utilization test.

### Serological confirmation of *Salmonella* isolates

The *Salmonella* isolates were also subjected to latex agglutination test using *Salmonella* Hi-latex identification kit (Hi-Media) for genus confirmation. The test was performed as per the directions provided by the manufacturer. The isolates showing visible agglutination within 2 min were considered positive.

### Molecular confirmation of *Salmonella* isolates

#### Isolation of genomic DNA

The DNA used for molecular studies was extracted using DNeasy blood and tissue kit (Qiagen, Germany) following the manufacturer’s instructions

#### Multiplex PCR for confirmation of genus and serovar

The PCR targeting *ompC* (*Salmonella* genus specific), *typh* (*S.* Typhimurium specific) genes of *Salmonella* was standardized as per the method described by **Alveraz *et al.* (2004)** with necessary modifications in composition of reaction mixture. The PCR reaction mixture for amplification consisted of 2.5 μl of 10x *Taq* buffer (Tris HCL with 15mM MgCl_2_), 1 μl of 2.5 mM concentration of each dNTPs, 1 μl (10 pmol) each of forward and reverse primers, 1 U *Taq* Polymerase, 5 μl of DNA template and nuclease-free water was added to make the final volume to 25μl.

The amplification was carried out in a thermal cycler. The cycling conditions used in mPCR consisted of an initial denaturation (95°C, 2 min), followed by 30 cycles of denaturation (95°C for 1 min), primer annealing (57 °C for 1 min) and extension (72°C, 2 min). A final extension at 72°C was given for 5 min.

The amplified PCR products were analysed using horizontal submarine gel electrophoresis in 2% agarose gel prepared in 0.5X TAE buffer and stained with 0.5 μg of ethidium bromide per ml. A separate well loaded with 100bp DNA ladder was run simultaneously. The amplified products were visualized and confirmed over a gel documentation system.

## Results

### Biochemical characterization of isolates

As many as 94 isolates of *Salmonella* procured from various north Indian states were used in the present study and checked for their morphology and biochemical characteristics. On BGA agar, colonies appeared as moderately large, moist, smooth and colourless with pink background. The isolates showing morphological characteristics identical to *Salmonella* were subjected to the several biochemical tests for their further identification. These test isolates exhibited characteristic reactions on TSI (Triple Sugar Iron) and Simmon’s citrate, whereas negative on urease agar slant confirming the isolates as *Salmonella* biochemically.

### Latex agglutination test

All the *Salmonella* isolates, showing putative morphological and biochemical characteristics were subjected to latex agglutination test for serological characterization. The isolates tested gave desired agglutination reaction confirming genus of the isolates (Fig. 1).

**Fig. 1:**
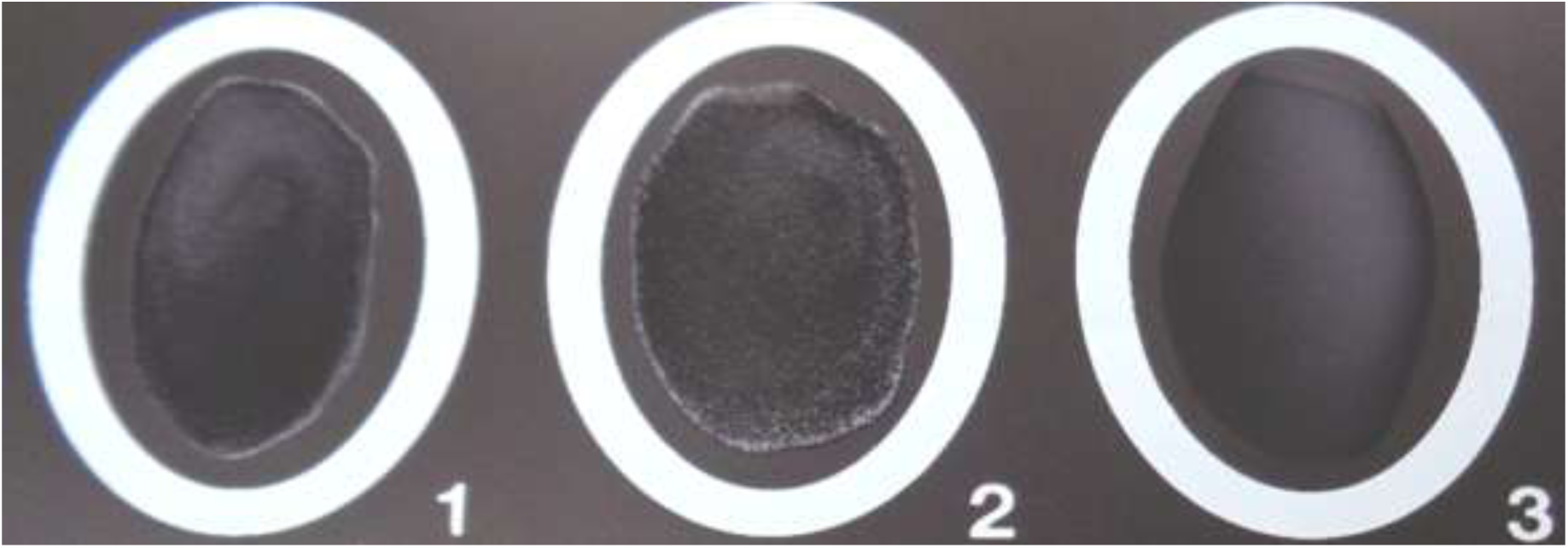
Latex Agglutination Test. (1- Positive control, 2- Positive isolate and 3- Negative control)

### Molecular confirmation of isolates

Conventional identification methods of *Salmonella* are based on culture growth using selective media and the biochemical characterization of the isolates. After confirmation using conventional identification methods, these isolates were subjected to highly sensitive, specific and reproducible molecular techniques, *viz.*, PCR for rapid detection and confirmation of *Salmonella* serotype. The genomic DNA of all *Salmonella* isolates were separated and subjected to the amplification with genus specific primer (*ompC*) and Typhimurium specific primer (*typh*) by mPCR. Multiplex PCR has an advantage of simultaneous detection of *Salmonella* genus and Typhimurium serovar. Optimized reaction mixture and cycling conditions for each gene revealed expected product size of 204 and 401 bp for *ompC* and *typh* genes, respectively (Fig. 2). The specificity of the primers was checked by using *S.* Typhimurium ATCC23564 as positive control and *E. coli* ATCC25922 as negative control. All the isolates yielded expected products, confirming both genus and serovar. Earlier workers have also reported the use of mPCR targeting *ompC* and *typh* genes for confirmation of *S.* Typhimurium (**Ahmed *et al.*, 2012; Anjay *et al.*, 2015**).

**Fig. 2:**
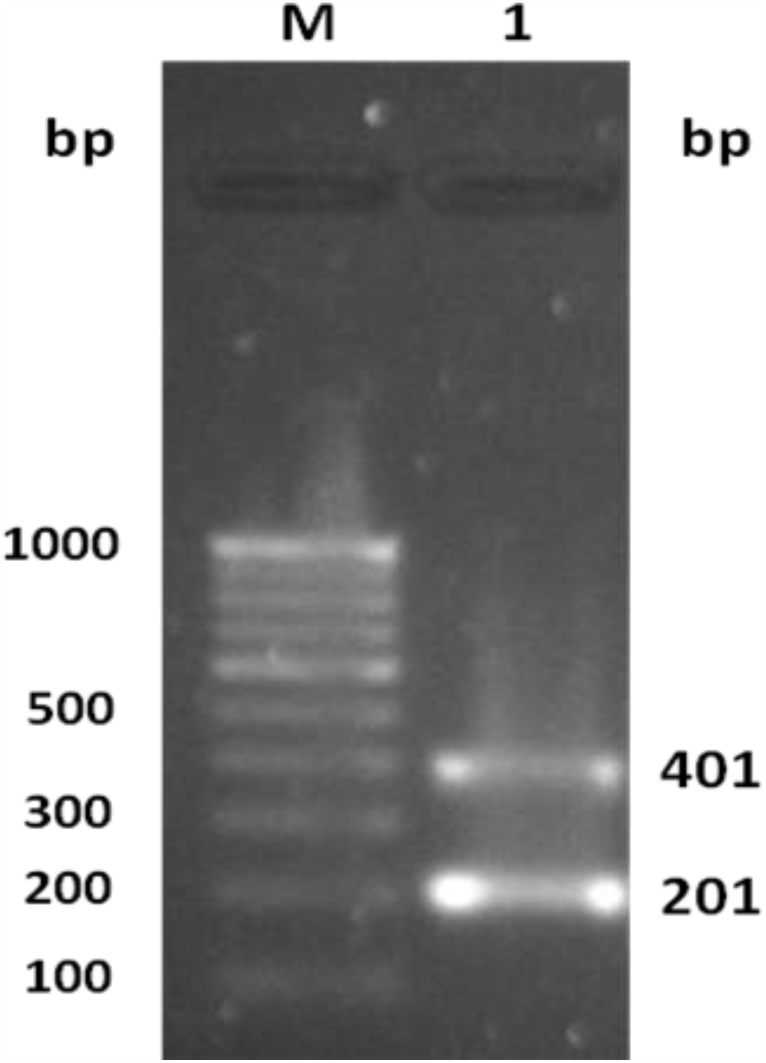
PCR product of *ompC* and *typh* genes. Lane M: 100 bp DNA ladder Lane 1: *ompC* and *typh* gene amplification in *Salmonella* Typhimurium isolate

All the 94 suspected *Salmonella* isolates of diverse origin from different parts of North Indian states were confirmed as *Salmonella* Typhimurium serovar by various biochemical, serological and molecular methods.

## Acknowledgment

This study was supported by a research grant “Outreach Programme on Zoonotic Disease” funded by Indian Council of Agriculture Research (ICAR), New Delhi. Authors acknowledge laboratory facilities extended by the Dean, College of Veterinary and Animal Sciences, G.B. Pant University of Agriculture and Technology, Pantnagar, Uttarakhand, India.

